# Red/near-infrared light activates the mitochondrial large-conductance calcium-activated potassium channel in glioblastoma cells

**DOI:** 10.64898/2026.04.02.716077

**Authors:** Piotr Bednarczyk, Joanna Lewandowska, Bogusz Kulawiak, Adam Szewczyk

## Abstract

Mitochondrial potassium channels, located in the inner mitochondrial membrane, play a crucial role in the cell’s life/death phenomenon. Activation of mitochondrial potassium channels by potassium channel openers may protect cells against ischemia-reperfusion injury. It is known that mitochondrial large conductance calcium-activated potassium channels interact with various mitochondrial proteins, including enzymes of the respiratory chain. Numerous studies indicate that the mitochondria, especially cytochrome c oxidase, play a crucial role as a chromatophore in the cellular response to red and near-infrared light. In this study, we employ the patch-clamp technique and single-channel recordings to investigate the regulation of glioblastoma mitochondrial large conductance calcium-activated potassium channel activity by infrared light. Specifically, we examined the effects of wavelengths 620 nm, 680 nm, 760 nm, and 820 nm in a redox-controlled environment. Our findings suggest that illuminating the inner mitochondrial membrane with these wavelengths may activate mitochondrial large conductance calcium-activated potassium channels. These results offer new insights into the regulation of mitochondrial potassium channels by cytochrome c oxidase, which may lead to the development of non-pharmacological interventions with potential cytoprotective benefits.

## INTRODUCTION

Mitochondria are central hubs of cellular bioenergetics, calcium signaling, redox regulation, and apoptosis, contributing to the metabolic flexibility of eukaryotic cells (Giorgi et al., 2018; Nicholls, 2016; Nunnari & Suomalainen, 2012; Spinelli & Haigis, 2018). Their activity depends on maintaining mitochondrial membrane potential (ΔΨm), oxidative phosphorylation (OXPHOS), and strict control of ion balance within the matrix and between the intermembrane space/cytosol. In recent years, mitochondrial ion channels have emerged as critical regulators of these processes, enabling fine-tuning of metabolism in response to cellular stress (O’Rourke, 2007; Szabo & Szewczyk, 2023). Among these, mitochondrial potassium (mitoK) channels are particularly important, as they control matrix volume, respiration, and the synthesis of reactive oxygen species (ROS) (Augustynek et al., 2017; Szabo & Szewczyk, 2023). In general, mitochondrial potassium channels play an important role in the cell life/death phenomenon and cell senescence (Garlid et al., 2003; Głuchowska et al., 2023; Leanza et al., 2012; Lewandowska et al., 2025; Szewczyk, 2024; Szewczyk et al., 2009; Szewczyk & Marbán, 1999). Several K⁺-conducting pathways have been described in the inner mitochondrial membrane (IMM), including ATP-sensitive (mitoK_ATP_), large-conductance Ca²⁺-activated (mitoBK_Ca_), voltage-dependent (mitoKv), and two-pore-domain channels such as TASK-3 (Bajgar et al., 2001; Szabò et al., 2005; Szabo & Szewczyk, 2023). Despite major progress, the molecular identity, regulation, and physiological roles of many mitochondrial K⁺ channels remain incompletely understood (Szewczyk et al., 2025).

Among mitochondrial potassium (mitoK) channels, the mitochondrial large-conductance calcium-activated potassium (mitoBK_Ca_) channel is of particular interest due to its role in cytoprotection (Dabrowska et al., 2022; Fretwell & Dickenson, 2009; Kampa et al., 2021; Xu et al., 2002). Activation of mitoBK_Ca_ channel causes mild mitochondrial depolarization, controlled matrix swelling, and reduced ROS synthesis during ischemia–reperfusion injury (Szteyn & Singh, 2020). The channel is modulated by matrix Ca²⁺ levels, the redox state, kinases, and interactions with respiratory chain complexes (Bednarczyk et al., 2013; Frankenreiter et al., 2017). Despite these findings, the upstream regulatory mechanisms that control mitoBK_Ca_ activation and could be harnessed for therapeutic stimulation remain unclear.

The mitoK channels interact with various mitochondrial proteins (Kathiresan et al., 2009; Peng et al., 2014; Sokolowski et al., 2011). Some of them are part of the respiratory chain. For instance, it has been suggested that mitochondrial ATP-sensitive potassium (mitoK_ATP_) channels interact with succinate dehydrogenase (Ardehali et al., 2004). Additionally, mitochondrial tandem pore domain potassium (mitoTASK-3) channels also interact with the respiratory chain (Yao et al., 2017). A recent report revealed a similar interaction between the mitoKv1.3 channel and the respiratory chain complex I (Peruzzo et al., 2020). In cardiac mitochondria, it was found that the β1 subunit of the mitoBK_Ca_ channels interacts with cytochrome oxidase (COX) subunit I (Ohya et al., 2005). Furthermore, studies have demonstrated that other respiratory chain protein complexes interact with mitoBK_Ca_ channels in both cardiac (J. Zhang et al., 2017), and brain mitochondria (Singh et al., 2016). This putative direct functional coupling between the energy-generating system (the respiratory chain) and the energy-dissipating system (potassium channels) may represent an important regulatory mechanism in mitochondria. In line with this idea, we found that the activity of mitoBK_Ca_ channels in glioblastoma U-87 MG cells is regulated by mitochondrial substrates and inhibitors acting through the respiratory chain (Bednarczyk et al., 2013). Taken together, these findings suggest that COX is a key regulator of these channels.

In parallel, interest has grown in photobiomodulation (PBM), the use of low-intensity red or near-infrared light (620 – 850 nm) to modulate cellular metabolism and survival pathways (Karu, 2010; W. Zhang et al., 2021). PBM enhances mitochondrial respiration, ATP synthesis, and redox balance, with protective or stimulatory outcomes depending on the biological context (Karu et al., 2004). Beneficial effects of PBM have been reported in neurodegeneration, cardiovascular diseases, and wound healing, suggesting that mitochondria contain light-sensitive components capable of driving adaptive metabolic responses (Abijo et al., 2023). However, the molecular mechanisms underlying light-driven mitochondrial regulation are still under investigation and appear to involve both primary photoreceptors and downstream ion-dependent modulation of bioenergetics.

Cytochrome c oxidase (complex IV, COX) is considered the main mitochondrial chromophore for red and near-infrared light (620, 680, 760, and 820 nm), mainly due to copper centers that undergo reversible redox transitions upon photon absorption (Karu, 2010; Karu et al., 2004). Light excitation of COX can increase enzymatic turnover, enhance electron transport, and improve oxygen utilization, ultimately boosting ATP generation under both physiological and stress conditions (Hamblin, 2018).

The mitoBK_Ca_ channel is particularly sensitive to redox modulation and closely coupled to the mitochondrial metabolic state (Lewandowska et al., 2024; Szabo & Szewczyk, 2023). Its open probability changes with thiol oxidation, ROS exposure, and interactions with respiratory chain components (Rotko et al., 2020). Moreover, mitoBK_Ca_ contributes to redox-dependent signaling during ischemic preconditioning, where transient oxidative shifts trigger mitochondrial protection (Goswami et al., 2019; Szteyn & Singh, 2020). These findings support the idea that PBM-induced COX redox changes might act as an upstream trigger modulating mitoBK_Ca_ function.

Despite growing evidence that PBM enhances mitochondrial performance and redox-sensitive ion transport, the mechanism by which light exposure, directly or indirectly, affects mitoBK_Ca_ activity remains unknown, as does whether such regulation depends on COX and the wavelength of illumination. No comprehensive study has determined if red light within the COX absorption spectrum can modulate mitoBK_Ca_ channel activity.

In this study, we investigated whether red and near-infrared light within the effective absorption range of cytochrome c oxidase (COX) modulates mitoBK_Ca_ channel activity. Using mitoplast preparations and patch-clamp recordings, we analyzed the channel’s light sensitivity under different redox conditions and across relevant wavelengths. We hypothesized that red light stimulates mitoBK_Ca_ *via* COX-dependent redox modulation, thereby influencing mitochondrial function. Our results reveal a novel connection between photobiomodulation, COX-driven redox signaling, and mitoBK_Ca_ channel activity, identifying this channel as a new component of light-responsive mitochondrial physiology. Specifically, we examined the effects of wavelengths 620, 680, 760, and 820 nm under controlled redox conditions. The data indicate, for the first time, that illumination of the inner mitochondrial membrane at specific wavelengths may activate mitoBK_Ca_ channels, uncovering a novel mechanism linking photobiomodulation to mitochondrial electrophysiology. These findings provide new insights into the regulation of mitoK channels and may contribute to the development of non-pharmacological interventions towards these proteins.

## MATERIAL AND METHODS

### Chemicals

Acrylamid, Bis–Tris, Coomassie blue G–Brillant Blau G250, glycerin, potassium hydroxide, and tricine were obtained from Carl Roth GmbH + Co., Germany. Digitonin and n-dodecyl-ß-D-maltoside were obtained from SERVA, Germany. Ammonium persulfate, potassium chloride, and tris(hydroxymethyl)aminomethane were purchased from BioShop, Canada. PBS–Dulbecco’s phosphate buffered saline w/o magnesium and w/o calcium was from Biowest, France. Protease inhibitor cocktail tablets complete EDTA-free were from Roche, Germany. RNase-free DNase set was from Qiagen AG, Germany. Tween-20 was purchased from Bio-Rad, USA. All other chemicals were from Sigma-Aldrich, USA.

### Cell culture

U-87 MG cells and the corresponding knockout cell lines were cultured in Dulbecco’s Modified Eagle Medium (Biowest) supplemented with 10% FBS (Gibco, USA), GlutaMAX (Gibco), 100 U/ml penicillin, and 100 µg/ml streptomycin (Sigma-Aldrich). Cells were maintained at 37 °C in a humidified atmosphere containing 5% CO₂ and passaged every 3–4 days. Cell identity was verified using short tandem repeat (STR) profiling as described before (Bednarczyk et al., 2013).

### Mitochondria preparation

Mitochondria were prepared from the human astrocytoma (glioblastoma) WT U-87 MG and ΔBK_Ca_ U-87 MG cells as previously described (Bednarczyk et al., 2013; Kampa et al., 2021, Kulawiak et al., 2023). Human astrocytoma cells from five culture flasks were collected in PBS medium and centrifuged at 800 × g for 10 min. The cell pellet was resuspended and homogenized in a preparation solution (250 mM sucrose, 5 mM HEPES, pH 7.2). For mitochondrial isolation, the homogenate was centrifuged at 9,200 × g for 10 min. The resulting pellet was resuspended and centrifuged at 780 × g for 10 min. The supernatant was transferred to a new tube and centrifuged at 9,200 × g for 10 min. Finally, the pelleted mitochondria were resuspended in a storage solution (150 mM KCl, 10 mM HEPES, pH 7.2) and centrifuged at 9,200 × g for 10 min. In the last step, the mitochondria were resuspended in approximately 0.3 ml of storage solution. All procedures were performed at 4°C.

### Patch-clamp experiments

Patch-clamp experiments were performed as previously described (Kampa et al., 2021; Maliszewska-Olejniczak et al., 2024; Walewska et al., 2022). In brief, mitoplasts were isolated from WT U-87 MG and ΔBK_Ca_ U-87 MG human astrocytoma mitochondria by placing them in a hypotonic solution (5 mM HEPES, 100 μM CaCl_2_, pH 7.2) for approximately 2 minutes to induce swelling and rupture of the outer membrane. A hypertonic solution (750 mM KCl, 30 mM HEPES, 100 μM CaCl_2_, pH 7.2) was added to restore isotonicity. The final bath isotonic solution contained 150 mM KCl, 10 mM HEPES, and 100 μM CaCl_2_ at pH 7.2. Similarly, the patch-clamp pipette was filled with an isotonic solution. Mitoplasts are easily identifiable by their size, rounded shape, transparency, and the presence of a “cap”, distinguishing them from cellular debris in the preparation. For all experiments, an isotonic solution containing 100 μM CaCl_2_ was used as the control. A low-calcium isotonic solution (1 μM CaCl_2_) contained 150 mM KCl, 10 mM HEPES, 1 mM EGTA, and 0.752 mM CaCl_2_ at pH 7.2. Channel modulators were added as dilutions in the isotonic solutions containing either 100 μM CaCl_2_ or 1 μM CaCl_2_. A perfusion system was used to apply these substances, consisting of a holder with a custom-made glass tube, a peristaltic pump, and Teflon tubing. Mitoplasts at the tip of the measuring pipette were transferred into the openings of a multibarrel “sewer pipe” system, where their outer faces were rinsed with the test solutions (Fig. 1A). All experiments were conducted in patch-clamp inside-out mode. The voltages reported correspond to those applied to the inside of the patch-clamp pipette, with positive potentials indicating the physiological polarization of the inner mitochondrial membrane (outside positive).

**Figure 1.**
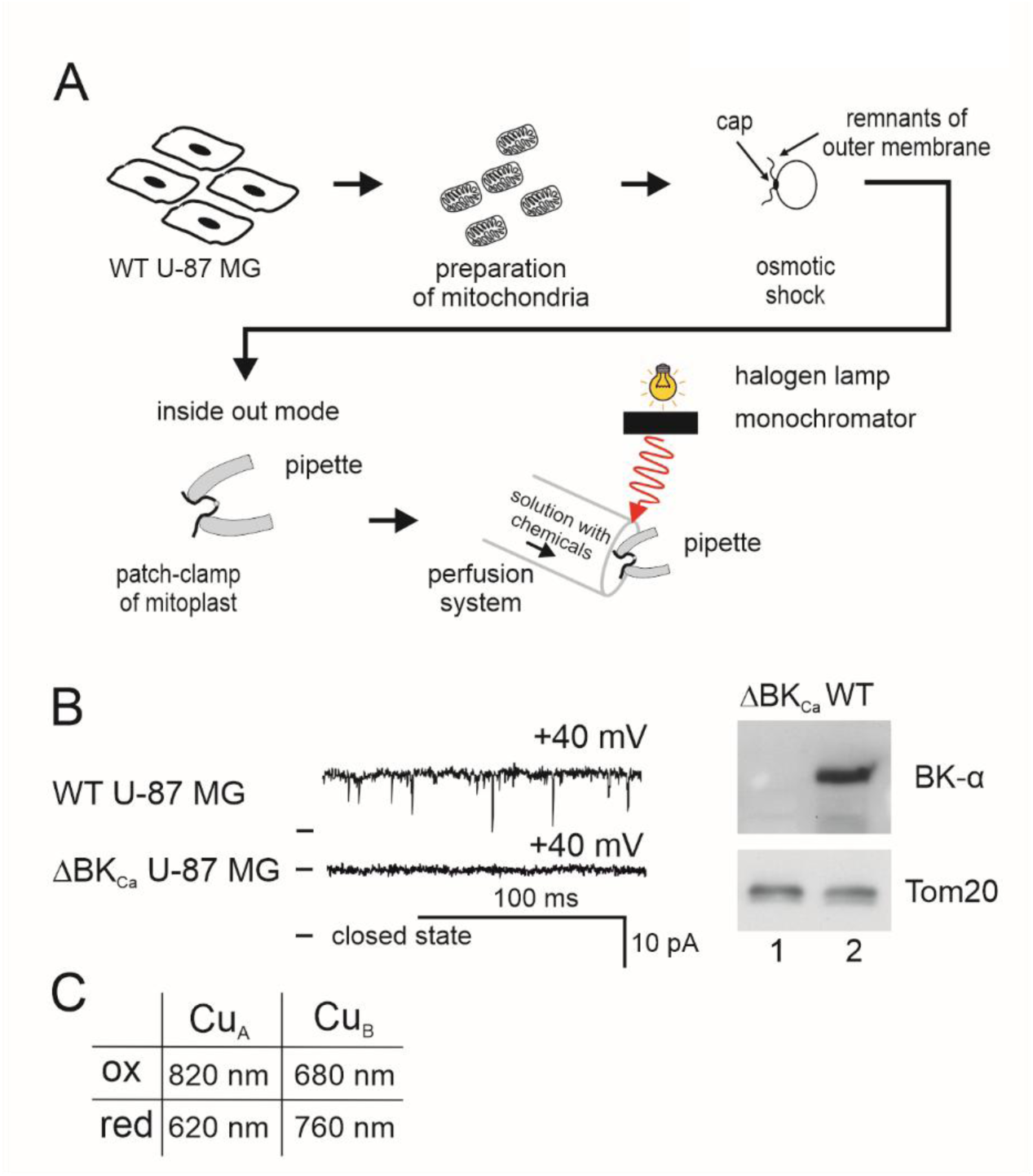
Single-channel recordings of the mitochondrial large-conductance calcium-regulated potassium channel in the presence of an illuminating system. **A.** Schematic representation of the mitoplast preparation, patching of the mitoplast, and, finally, the patch-clamp experiment in the inside-out mode by means of illumination. The experiments were carried out in patch-clamp inside-out mode. The matrix side of the mitochondrial membrane is exposed to externally added substances. **B.** Single-channel recordings of the mitoBK_Ca_ channel in the inner mitochondrial membrane of WT U-87 MG cells, and recording from ΔBK_Ca_ U-87 MG cells (left panel). Immunoblot of astrocytoma mitochondria from WT U-87 MG, and ΔBK_Ca_ U-87 MG cells labeled with the anti-BK_Ca_ channel a subunit and Tom20 antibody (right panel). **C.** Redox state and red/near-infrared light wavelengths used in the experiments.

The electrical connection was established using Ag/AgCl electrodes and an agar salt bridge (3 M KCl) as the ground electrode. The current was recorded with a patch-clamp amplifier (Axopatch 200B, Molecular Devices, USA). Pipettes, made of borosilicate glass, had a resistance of 10–20 MΩ and were pulled using a Narishige PC-10 puller. The current was low-pass filtered at 1 kHz and sampled at 100 kHz. Single-channel recordings were analyzed, with channel conductance calculated from the current–voltage relationship. The probability of channel opening (Po, channel open probability) was determined using Clampfit software. The single-channel search mode was used for analysis. Results are presented as mean values ± standard deviation (S.D.). In figures showing single-channel recordings, “-” indicates the closed state of the channel.

### Illumination protocol

The illumination set-up contains: 150 W halogen lamp, MSH-150 monochromator systems, and glass fiber bundles (380 – 1600 nm) (Quantum Design). The beam was positioned on the patch-clamp pipette at 550 nm (green) (Fig. 1A). The final power densities at 620 nm, 680 nm, 760 nm, and 820 nm were equal to 510, 470, 430, and 335 μW/cm^2^, respectively. The light power densities were measured using a Microscope Slide Power Meter Sensor Head S170C (Thorlabs).

### SDS-PAGE and immunoblotting

WT U-87 MG and ΔBK_Ca_ U-87 MG cells were seeded in 6-well plates, washed with cold PBS, and detached from the wells. Cells were collected by centrifugation and lysed in RIPA buffer supplemented with protease inhibitors. After incubation on ice, the lysates were clarified by centrifugation at 4 °C. The resulting supernatants were collected and stored at −20 °C for further analysis. Aliquots were taken for protein concentration determination prior to electrophoretic analysis. Equal amounts of whole-cell lysates solubilized in Laemmli buffer (Bio-Rad) were separated by 10% Tris–tricine SDS-PAGE and subsequently transferred onto polyvinylidene difluoride (PVDF) membranes (Bio-Rad). After transfer, membranes were blocked with 10% non-fat dry milk prepared in Tris-buffered saline containing Tween 20 and incubated with primary antibodies: anti-BK_Ca_ (NeuroMabs, USA, clone L6/60, 1:200) and anti-Tom20 (Sigma, 1:1000, F118569). Immunodetection was performed using HRP-conjugated secondary antibodies against rabbit or mouse IgG (GE Healthcare and Thermo Fisher Scientific) followed by visualization with enhanced chemiluminescence reagents (GE Healthcare). Molecular weight estimation was carried out using the PageRuler Prestained Protein Ladder (Thermo Fisher Scientific).

### Statistical analysis

All experiments were performed with at least three or more independent biological replicates to ensure reproducibility. Results are presented as mean ± S.D., as calculated using Prism 4 (GraphPad Software Inc.). One-way ANOVA was employed to analyze the experimental data. P-values were considered significant as follows: *p ≤ 0.05, **p ≤ 0.01, and ***p ≤ 0.001.

## RESULTS

### Biophysical and pharmacological properties of the mitoBK_Ca_ channels in astrocytoma

To characterize the biophysical and pharmacological properties of the mitochondrial large-conductance calcium-activated potassium (mitoBK_Ca_) channel, the patch-clamp technique was used. Experiments were performed on mitoplasts isolated from human astrocytoma WT U-87 MG and ΔBK_Ca_ U-87 MG cells. A schematic diagram illustrating mitoplast preparation, patching, and the inside-out patch-clamp configuration with illumination is shown in Fig. 1A. For confirmation of the channel presence, single-channel recordings of the mitoBK_Ca_ channel in the inner mitochondrial membrane of WT U-87 MG cells were conducted (Fig. 1B). As negative control we used a cell line that we had previously described, in which the KCNMA1 gene encoding the pore-forming α subunit of the BK_Ca_ channel had been disrupted using the CRISPR/Cas9 technique (Kulawiak et al., 2023). The current was measured in a symmetric 150/150 mM KCl isotonic solution containing 100 µM CaCl_2_. Electrophysiological experiments demonstrated that the disruption of the *KCNMA1* gene resulted in the loss of BK_Ca_-type channels (Fig. 1B, left panel). This observation was confirmed by immunoblot analysis of astrocytoma mitochondria from WT U-87 MG and ΔBK_Ca_ U-87 MG cells labeled with the anti-BK_Ca_ channel α subunit and Tom20 antibodies (Fig. 1B, right panel).

In 53 patches, we observed a single-channel conductance of 286 ± 3 pS, calculated from the mean of the current-voltage relationship (Fig. 2A). The current showed no signs of rectification. The channel open probability (Po) increased from ∼approximately 0.11 at −80 mV to ∼approximately 0.97 at positive voltages, representing the typical voltage dependence for mitoBK_Ca_ channels (Fig. 2B). Representative single-channel recordings for the mitoBK_Ca_ channel measured at different voltages from + 40 mV to −40 mV in a symmetrical isotonic solution (150 mM KCl, 10 mM HEPES, and 100 μM CaCl_2_, pH 7.2) was shown in figure 2C. To test the calcium sensitivity of the channel, we reduced the Ca^2+^ concentration from high (100 µM) to low (1 µM) (Fig. 2D). Analysis of channel open probability (Po) revealed a statistically significant inhibition by 1 μM Ca^2+^. After adding the BK_Ca_ channel opener NS21021, Po increased from 0.08 to 0.34 (Fig. 2D). In the next step, to confirm that the observed channel is of the BK_Ca_ type, we used inhibitors such as paxilline. Figure 2E demonstrates the inhibitory effect of paxilline. The control channel open probability decreased from ∼97.8% to ∼0.1% in the presence of 1 μM paxilline.

**Figure 2.**
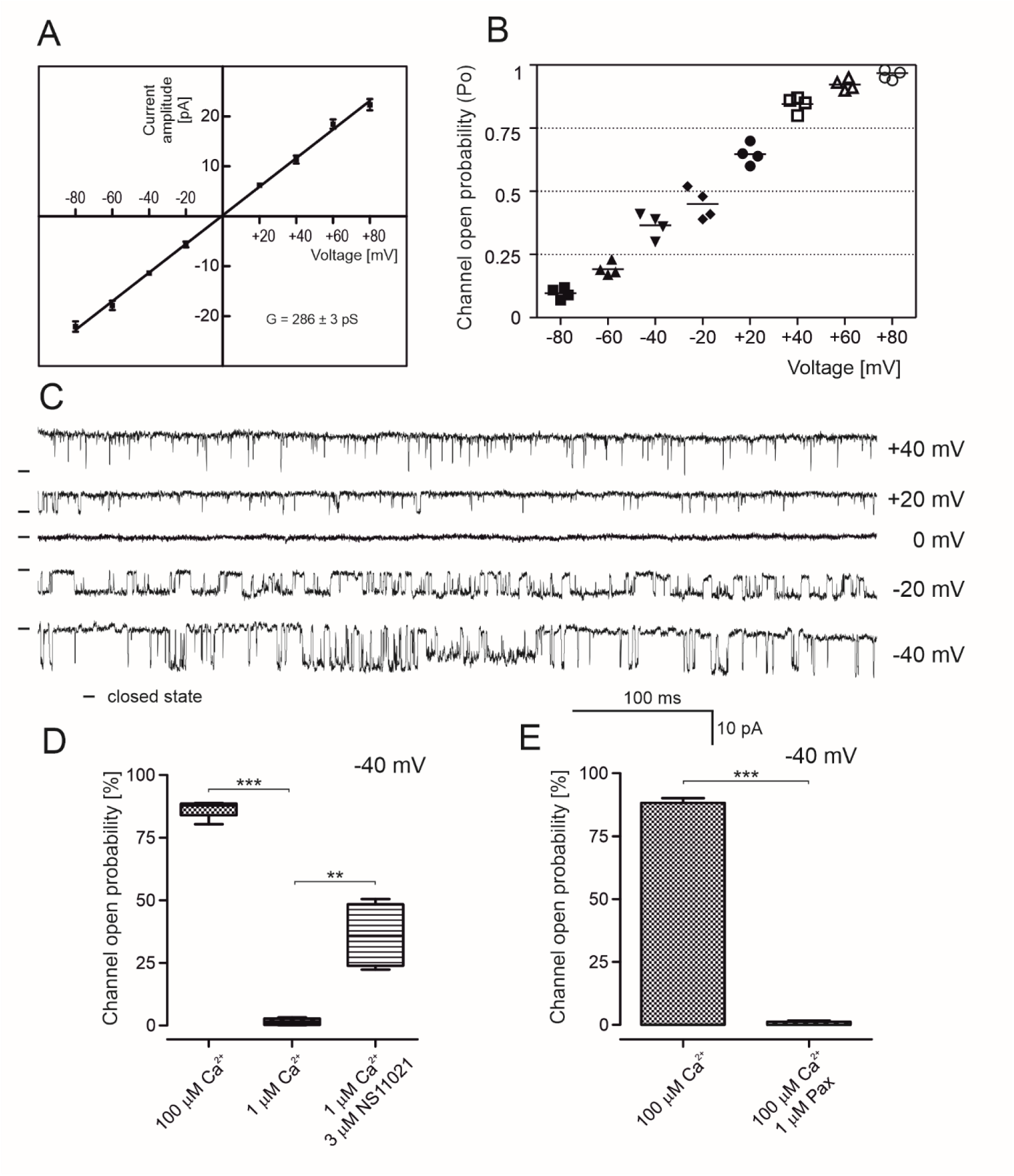
Electrophysiological and pharmacological properties of the mitoBK_Ca_ channel in U-87 MG mitochondria. **A.** Current-voltage (i-v) characteristics of single-channel events. The conductance was established at 286 ± 3 pS (n = 4). **B.** Channel open probability (Po) of the mitoBK_Ca_ under control conditions at different voltages (n = 4). **C.** Representative single-channel recordings in symmetric 150/150 mM KCl isotonic solution with a high calcium concentration (100 µM Ca^2+^) at different voltages. **D.** Channel open probability analysis of the single-channel recordings in the presence of 100 mM Ca^2+^, 1 mM Ca^2+^, and 3 mM NS11021. The effect is reversible upon the addition of 3 µM NS11021 (n= 3). ***P<0.001 vs. the control. **P<0.001 vs. 1 µM Ca^2+^. **E.** Effects of 1 µM paxilline (Pax) on the single-channel activity. The distribution of the probability of channel opening under the above conditions is shown with ***P<0.001 vs. the control (n = 4).

### Light regulation of the mitoBK_Ca_ channels activity

Numerous studies have demonstrated that mitochondria, particularly cytochrome c oxidase, serve as key chromophores in cellular responses to red and near-infrared light (Desmet et al., 2006; Sanderson et al., 2018). The stimulation wavelength is shown in Figure 1C. In this study, we employed patch-clamp techniques and single-channel recordings to examine how the redox state of the respiratory chain and red/near-infrared light regulate the activity of mitoBK_Ca_ channels in glioblastoma cells. Specifically, we investigated the effects of illumination at 620, 680, 760, and 820 nm. In the first step, we have investigated how oxidized respiratory chain state (in the presence of 0.3 mM ferricyanide K_3_Fe(CN)_6_) and red/near-infrared illumination modulate the activity of mitoBK_Ca_ channels. Ferricyanide acts as an artificial electron acceptor in mitochondria and it accepts electrons from components on the respiratory chain specifically at the level of cytochrome c or cytochrome bc1. In Figure 3A, a schematic illustrates the redox sensitivity of the Cu_A_ center, showing that the reduced state preferentially absorbs at 620 nm, whereas the oxidized state exhibits an absorption peak around 820 nm. Quantification of channel open probability shows that, under control conditions, channels exhibit high activity at +40 mV and moderate activity at −40 mV. Application of the oxidizing agent K_3_Fe(CN)_6_ (0.3 mM) markedly suppressed channel activity at both voltages (Fig. 3B). Illumination at 620 nm did not restore activity under oxidizing conditions, and the differences relative to control were highly significant.

**Figure 3.**
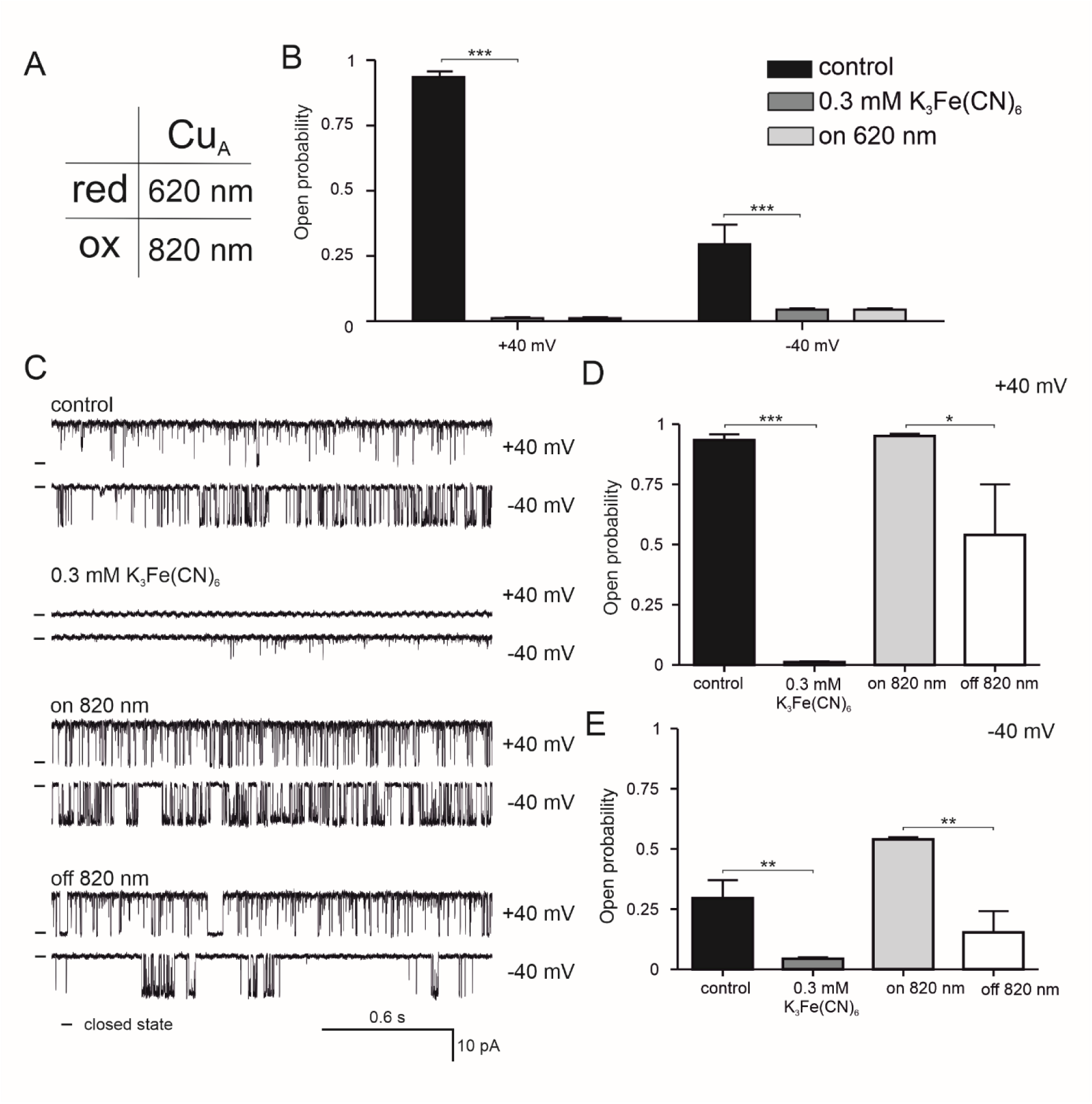
The mitoBK_Ca_ channel activity measurements and red/infrared illumination under the oxidative conditions. **A.** Wavelengths 620 nm and 820 nm were used in the experiments. **B.** Channel open probability (Po) of the mitoBK_Ca_ under control, in the presence of 0.3 mM K_3_Fe(CN)_6_, and after illumination with 620 nm at +40 mV, and -40 mV. **C.** Single-channel recordings in symmetric 150/150 mM KCl isotonic solution with a high calcium concentration (100 µM Ca^2+^) at +40 mV, and -40 mV under control, in the presence of 0.3 mM K_3_Fe(CN)_6_, after illumination with 820 nm, and after switching off. **D.** and **E.** Channel open probability (Po) of the mitoBK_Ca_ under condition in the panel C. Data were presented with SD. Statistical significance of measurements compared to controls was determined at ***P<0.001, **P<0.01, and *P<0.05 (***) by one-way ANOVA.

A markedly different response was observed upon illumination at 820 nm. Representative single-channel recordings demonstrated these effects at +40 mV and −40 mV (Fig. 3C). Similar to previous observations, robust channel openings were observed in control conditions, whereas oxidizing conditions significantly reduce activity. Illumination at 820 nm restored frequent channel openings, whereas turning the light off again reduced the channel’s activity. At +40 mV, the open probability of the mitoBK_Ca_ was near maximal in control conditions and was almost completely abolished by K_3_Fe(CN)_6_. Illumination at 820 nm significantly rescued channel activity, which partially declined after the light is switched off (Fig. 3D). At −40 mV, a similar pattern was observed: oxidizing conditions significantly reduced the open probability, whereas illumination at 820 nm significantly increased channel activity compared with both the control and oxidized states. Removal of illumination again decreased overall activity (Fig. 3E). In general, the data indicate that mitochondrial channel activity is strongly suppressed by respiratory chain oxidation but can be reversibly enhanced by near-infrared light at 820 nm, consistent with the redox-dependent modulation of mitochondrial function.

In the next step, we demonstrated that modulation similar to that of the Cu_B_ redox center strongly influences mitoBK_Ca_ channel activity, and that this effect is wavelength-dependent. As summarized in Figure 4A, the oxidized and reduced states of Cu_B_ correspond to preferential absorption at 680 nm and 760 nm, respectively. At +40 mV, channels displayed a high probability of opening under control conditions (Fig. 4B). Chemical reduction with 0.5 mM TMPD/ascorbate markedly suppressed channel activity, reducing the open probability to nearly zero. Illumination at 680 nm did not reverse this inhibition, and channel activity remained low both during and after illumination (Fig. 4B), indicating that this wavelength does not effectively modulate channel gating under reducing conditions. In contrast, illumination at 760 nm robustly restored channel activity in the presence of TMPD/ascorbate. Representative single-channel recordings (Fig. 4C) show frequent and prolonged channel openings during 760 nm illumination, whereas activity was minimal in reducing conditions without light. Quantitative analysis confirmed a significant increase in open probability upon illumination at 760 nm compared with TMPD/ascorbate alone (Fig. 4D). Turning off the light led to a partial but significant decline in activity, indicating a reversible, light-dependent effect. Together, these data demonstrate that reduction suppresses mitoBK_Ca_ channel opening, but near-infrared light at 760 nm selectively counteracts this inhibition, restoring channel activity. This supports a model in which mitochondrial channel gating is regulated by the redox state, similar to the Cu_B_, and can be dynamically modulated by wavelength-specific photostimulation.

**Figure 4.**
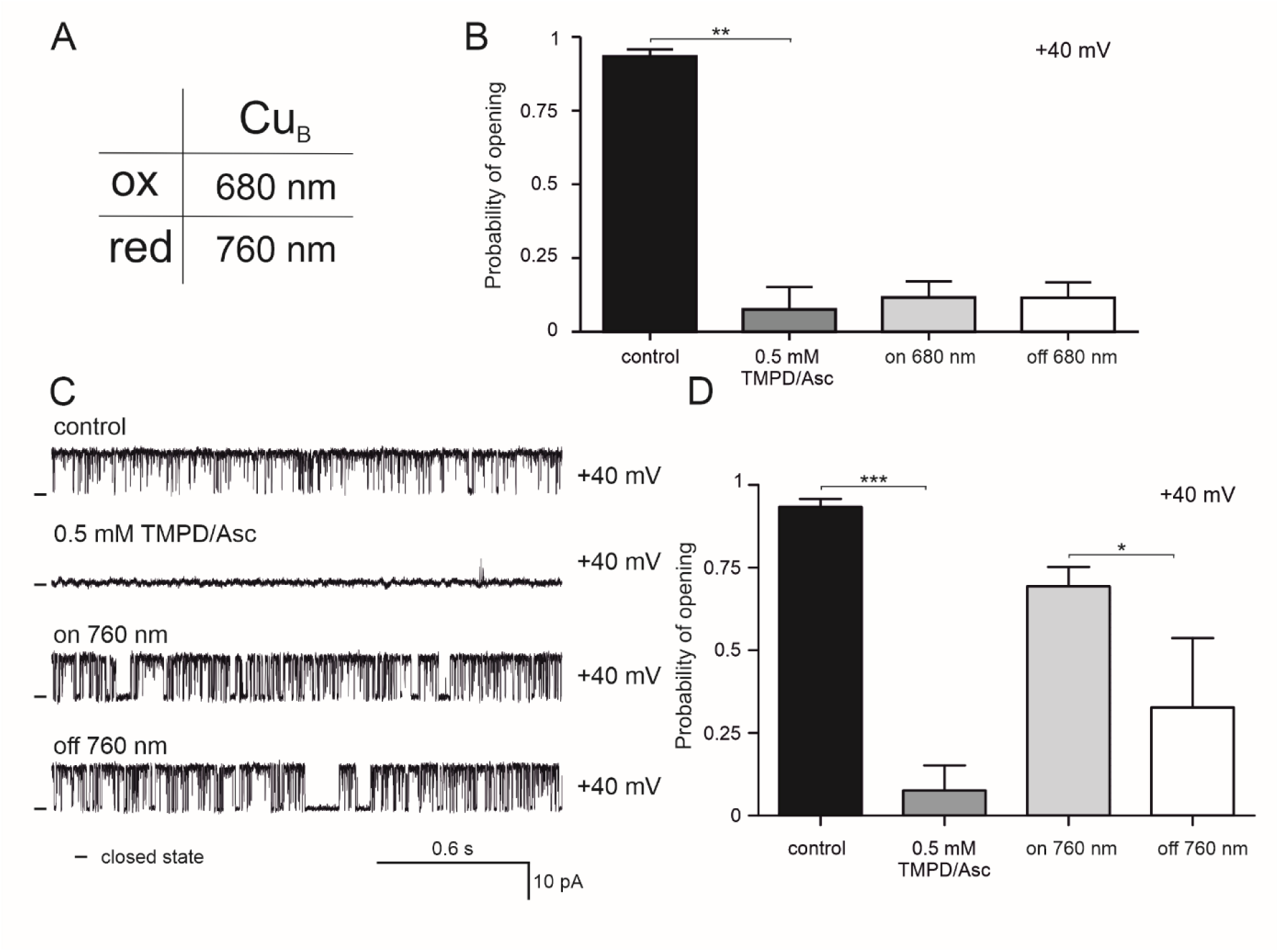
The mitoBK_Ca_ channel activity measurements and red/infrared illumination under the reduced conditions. **A.** Wavelengths 680 nm and 760 nm were used in the experiments. **B.** Open probability (Po) of the mitoBK_Ca_ channel under control, in the presence of 0.5 mM TMPD/Asc, and after illumination with 680 nm, and after switching off. Data were shown at +40 mV and -40 mV. **C.** Single-channel recordings in symmetric 150/150 mM KCl isotonic solution with a high calcium concentration (100 µM Ca^2+^) at +40 mV under control, in the presence of 0.5 mM TMPD/Asc, after illumination with 760 nm, and after switching off. **D.** Channel open probability (Po) of the mitoBK_Ca_ under conditions in panel C. Data were presented with SD. Statistical significance of measurements compared to controls was determined at ***P<0.001, **P<0.01, and *P<0.05 by one-way ANOVA.

In Figure 5, we summarize the dependence of mitoBK_Ca_ channel gating on the redox state. At +40 mV, channel open probability was high under control conditions, whereas both oxidative and reductive redox manipulations markedly suppressed channel activity (Fig. 5A). Oxidation reduced the channel open probability to near zero, while reduction also significantly decreased channel opening compared with control, indicating that maximal channel activity requires an intermediate redox state. Despite these pronounced effects on gating, single-channel current amplitude remained unchanged (Fig. 5B). Illumination at 820 nm (targeting Cu_A_ under oxidizing conditions) or 760 nm (targeting Cu_B_ under reducing conditions) did not significantly alter unitary current amplitude, indicating that light and redox modulation primarily affect channel open probability rather than conductance. Representative single-channel recordings further illustrate the wavelength- and redox-specific effects (Fig. 5C–F). For Cu_B_, control conditions exhibited frequent channel openings that were unaffected by 820 nm illumination (Fig. 5C). In contrast, oxidation with 0.3 mM K_3_Fe(CN)_6_ strongly suppressed channel activity, and neither 760 nm nor 680 nm illumination restored openings (Fig. 5D).

**Figure 5.**
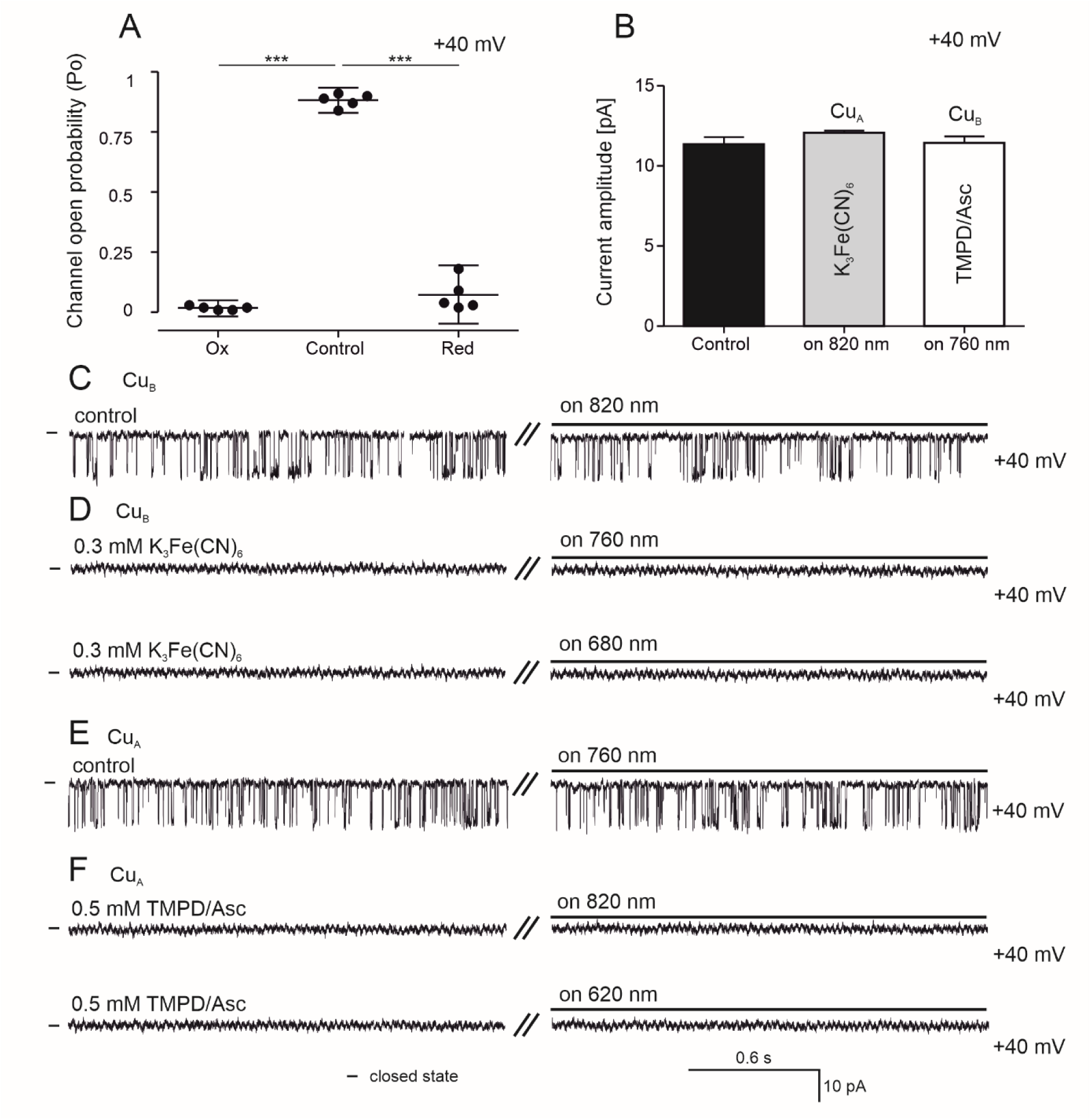
The mitoBK_Ca_ channel activity measurements and red/infrared illumination. **A.** Analysis of the mitoBK_Ca_ channel open probability in Ox and Red conditions at +40 mV. **B.** Current amplitude measured in the presence of 0.3 mM K_3_Fe(CN)_6_, and 0.5 mM TMPD/Asc after illumination with 820 nm and 760 nm, respectively. **C.** Single-channel recordings in symmetric 150/150 mM KCl isotonic solution with a high calcium concentration (100 µM Ca^2+^) (+40 mV) at control, and after illumination with 820 nm. **D.** Single-channel recordings in symmetric 150/150 mM KCl isotonic solution with a high calcium concentration (100 µM Ca^2+^) at +40 mV in the presence of 0.3 mM K_3_Fe(CN)_6_, and after illumination with 760 nm (upper panel) and 680 nm (bottom panel). **E.** Single-channel recordings in symmetric 150/150 mM KCl isotonic solution with a high calcium concentration (100 µM Ca^2+^) (+40 mV) at the control and after illumination with 760 nm. **F.** Single-channel recordings in symmetric 150/150 mM KCl isotonic solution with a high calcium concentration (100 µM Ca^2+^) at +40 mV in the presence of 0.5 mM TMPD/Asc, and after illumination with 820 nm (upper panel) and 620 nm (bottom panel). Data were presented with SD. Statistical significance of measurements compared to controls was determined at ***P<0.001 by one-way ANOVA.

For CuA, channels were active under control conditions and remained active during 760 nm illumination (Fig. 5E). However, reduction with 0.5 mM TMPD/ascorbate nearly abolished channel activity, and illumination at either 820 nm or 620 nm failed to rescue channel opening (Fig. 5F).

Together, these results demonstrate that mitochondrial channel gating is tightly regulated by the redox state of cytochrome c oxidase copper centers. While specific wavelengths can modulate channel activity in defined redox conditions, neither oxidation nor reduction alone supports high channel open probability, and light does not affect single-channel conductance.

## DISCUSSION

Our previous findings demonstrated that the activity of the mitoBK_Ca_ channel is modulated by the redox state of the respiratory chain, suggesting a close functional and possibly structural coupling of the mitochondrial potassium channel with cytochrome c oxidase (COX) in the inner mitochondrial membrane of human astrocytoma U-87 MG cells (Bednarczyk et al., 2013). Since the COX complex absorbs red/near-infrared light, we investigated whether this absorption affects the activity of the mitoBK_Ca_ channel.

### Infrared light stimulates mitoBK_Ca_ channel activity

In the present study, we demonstrated that light in the red and near-infrared spectral range, absorbed by COX, could modulate mitoBK_Ca_ channel activity. Cytochrome c oxidase is considered the main mitochondrial chromophore for red and near-infrared light (600-850 nm) due to its heme a, heme a3/Cu_B_, and Cu_A_/Cu_B_ centers that undergo reversible redox transitions upon photon absorption (Dyuba et al., 2013; Farver et al., 2006; Mason et al., 2009, 2014; Szundi et al., 2001). Red/near-infrared light absorption by these centers, particularly the Cu_A_/Cu_B_ centers, enhances the enzyme’s catalytic efficiency in reducing oxygen, thereby in fact increasing ATP synthesis (Jagdeo et al., 2012). This stimulation of COX activity initiates downstream events, including altered levels of ROS and nitric oxide, which are crucial for intracellular signaling (Pope & Denton, 2023). While the exact mechanisms remain under investigation, several studies suggest that light-induced COX activation influences mitochondrial membrane potential (Sanderson et al., 2018). This kind of photobiomodulation effect, wherein red/near-infrared light stimulates biological processes, is thought to promote glycolysis and ATP synthesis by enhancing electron transport through the mitochondrial respiratory chain (De Freitas & Hamblin, 2016; Pope & Denton, 2023). This activation of COX may further lead to alterations in intracellular calcium ion levels (Maghfour et al., 2024). Finally, these modulations activate various signaling pathways, which then contribute to downstream effects on cellular proliferation, migration, and differentiation, probably leading to therapeutic outcomes such as reduced inflammation and enhanced tissue repair (De Freitas & Hamblin, 2016; Hamblin, 2018; Maghfour et al., 2024).

Our current observation provides novel evidence that light energy, through COX-dependent redox transitions, may influence the gating behavior of the mitoBK_Ca_ channel. The experimental observations reported here can be summarized as follows. We demonstrated, using electrophysiological techniques, that mitoBK_Ca_ channels in glioblastoma cells can be modulated by specific wavelengths of red/near-infrared light (620, 680, 760, and 820 nm) under redox-controlled conditions. Using patch-clamp and single-channel recordings, we observed that illumination of the inner mitochondrial membrane enhances mitoBK_Ca_ activity. Since COX is a key photoacceptor in the red/near-infrared wavelength range, our results indicate that its light-induced redox modulation can translate into functional changes in mitoBK_Ca_ channel activity, linking photobiomodulation to mitochondrial ion channel regulation. The light sensitivity of mitoBK_Ca_ reveals that mitochondria can use light as an additional signal to regulate their metabolism and redox state.

This finding supports the concept that the mitoBK_Ca_ channel operates as a redox- and light-sensitive effector of mitochondrial activity. Through its ability to modulate potassium flux and, consequently, matrix volume and membrane potential, the channel may serve as a rapid feedback element coupling respiratory activity to ion conductance via the inner mitochondrial membrane. The involvement of COX as both an electron-transfer enzyme and a photoacceptor introduces a unique mechanism by which light can indirectly regulate mitochondrial bioenergetics via modulation of potassium channels. This dual sensitivity could enable mitochondria to integrate chemical and light signals, expanding their adaptive capacity under varying physiological or environmental conditions.

From a physiological perspective, the light-induced activation of mitoBK_Ca_ channels may have cytoprotective implications similar to those observed with pharmacological channel openers (Szewczyk, 2024). Previous studies have shown that activation of the mitoBK_Ca_ channel reduces mitochondrial calcium overload, limits ROS generation, and enhances cellular survival under stress conditions (Rotko et al., 2020). Might modulation of mitoBK_Ca_ by specific wavelengths absorbed by COX provide a non-pharmacological approach to achieving similar protective effects in cardiac and brain tissue? This mechanism could partly explain the beneficial effects of red and near-infrared light therapies reported in various models of neuroprotection, ischemic injury, and tissue repair.

Ultimately, our findings offer new insights into the biophysical basis of photobiomodulation. They suggest that mitochondrial ion channels, long recognized as key regulators of energy metabolism and redox balance, may also act as photoreactive components of the organelle (Szabo and Szewczyk, 2023). Future work should aim to elucidate the molecular interface between COX and mitoBK_Ca_, identify potential cofactors that mediate this light coupling, and determine whether similar mechanisms operate in other mitochondrial potassium channels, such as mitoK_ATP_ or mitoK_v_ channels. Understanding these interactions may contribute to the rational design of light-based interventions targeting mitochondrial function in various tissues.

### A possible mechanism of regulation of mitochondrial potassium channels by near-infrared irradiation absorbed by cytochrome c oxidase

How does red/near-infrared light absorbed by cytochrome c oxidase (COX) influence the activity of mitoBK_Ca_ channels? The mechanism underlying this phenomenon remains difficult to explain. It likely involves redox-dependent processes, gasotransmitter signaling, or potential structural coupling between COX and mitoBK_Ca_ channel.

Cytochrome c oxidase (COX) is the terminal enzyme of the mitochondrial electron transport chain (ETC), responsible for the reduction of molecular oxygen to water. Owing to its copper centers (Cu_A_, Cu_B_) and heme groups (heme *a* and *a_3_*) COX is considered the primary photoacceptor of NIR light. Absorption of photons at wavelengths such as 760 nm and 820 nm induces changes in the redox state of the enzyme. In its oxidized form, NIR irradiation promotes dissociation of nitric oxide (NO) from COX, leading to enhanced ETC activity, increased ATP synthesis, and elevated synthesis of reactive oxygen species (ROS) (Hamblin 2018). These could influence the mitochondrial membrane potential (ΔΨm) and redox balance, thereby indirectly modulating ion channel activity.

Mitochondrial potassium channels, including mitoBK_Ca_ orf mitoK_ATP_, are regulated by the mitochondrial redox state, ROS levels, gasotransmitters (NO, H_2_S, CO), and protein–protein interactions, including potential associations with COX (Bednarczyk et al., 2013). Respiratory substrates modulate mitoBK_Ca_ activity, suggesting functional coupling between the channel and COX, which may act as a redox sensor transmitting signals to K^+^ channels (Bednarczyk et al., 2013). In our paper we have shown that NIR irradiation at 760 or 820 nm activates mitoBK_Ca_ in a manner dependent on the redox state of the ETC; under oxidized conditions, NIR increases K^+^ current. These observations support the concept that COX is probably a key regulatory element that absorbs NIR light and modulates channel activity through redox-dependent mechanisms.

The proposed regulatory mechanisms can be grouped into three pathways. The first one concerns a direct redox/gasotransmitter pathway. NIR absorption by COX leads to NO dissociation from the catalytic heme *a*3–CuB center, resulting in increased ETC activity and production of ATP and moderate levels of ROS e.g., H_2_0_2_, O^2-^ (Hamblin 2018). ROS can modify cysteine residues in potassium channels (e.g., sulfenylation, S-glutathionylation, or S-sulfhydration), thereby activating mitoBKCa channel. For instance, H202 activates mitoKATP under oxidative stress, preventing opening of the mitochondrial permeability transition pore (mPTP). In addition, NO produced by COX acting as a nitrite reductase under NIR irradiation may directly activate channels via S-nitrosylation or indirectly through redox-sensitive kinases such as PKC or PKG (Kashiwagi et al., 2023).

Second pathway may concern the indirect mitochondrial signaling i.e. NIR-induced changes in ΔΨm and the proton gradient can modulate voltage-sensitive channels such as mitoBKCa channel. In the context of ischemia/reperfusion injury, activation of mitochondrial K^+^ channels by NIR via COX reduces excessive ROS production and enhances cytoprotection.

Third pathway may be based on direct structural interactions of COX and channel protein. In cardiac and brain mitochondria, the pore forming and regulatory subunits of mitoBKCa physically interacts with subunits of COX, as demonstrated by blue native electrophoresis, co-_immunoprecipitation_ experiments and other methods (Ohya et al., 2005; Bednarczyk et al., 2013; Singh et al., 2016; Zhang et al., 2017). Similar structural associations have been reported for other mitochondrial potassium channels, such as mitoK_ATP_ with succinate dehydrogenase (complex II) and mitoKv1.3 with complex I of the ETC (Ardehali et al., 2004; Peruzzo et al., 2020). This mechanism remains hypothetical and requires further investigation at the molecular level, including high-resolution structural approaches such as cryo-electron microscopy to elucidate protein–protein interactions.

### Final remarks

In summary, our results identify the mitoBK_Ca_ channel as a novel target of photobiomodulation mediated by cytochrome c oxidase (Figure 6). This finding expands the current understanding of how mitochondrial function can be modulated by red/infra-red light, revealing a previously unrecognized pathway linking COX photoactivation to mitochondrial potassium conductance. By demonstrating that specific wavelengths within the red and near-infrared range can alter mitoBK_Ca_ channel activity, our study bridges two major fields of mitochondrial research - bioenergetic regulation and photobiology. These results not only provide mechanistic insight into light-dependent mitochondrial responses but also highlight the mitoBK_Ca_ channel as a potential effector contributing to the beneficial outcomes of red-light exposure observed in numerous experimental and clinical settings.

**Figure 6.**
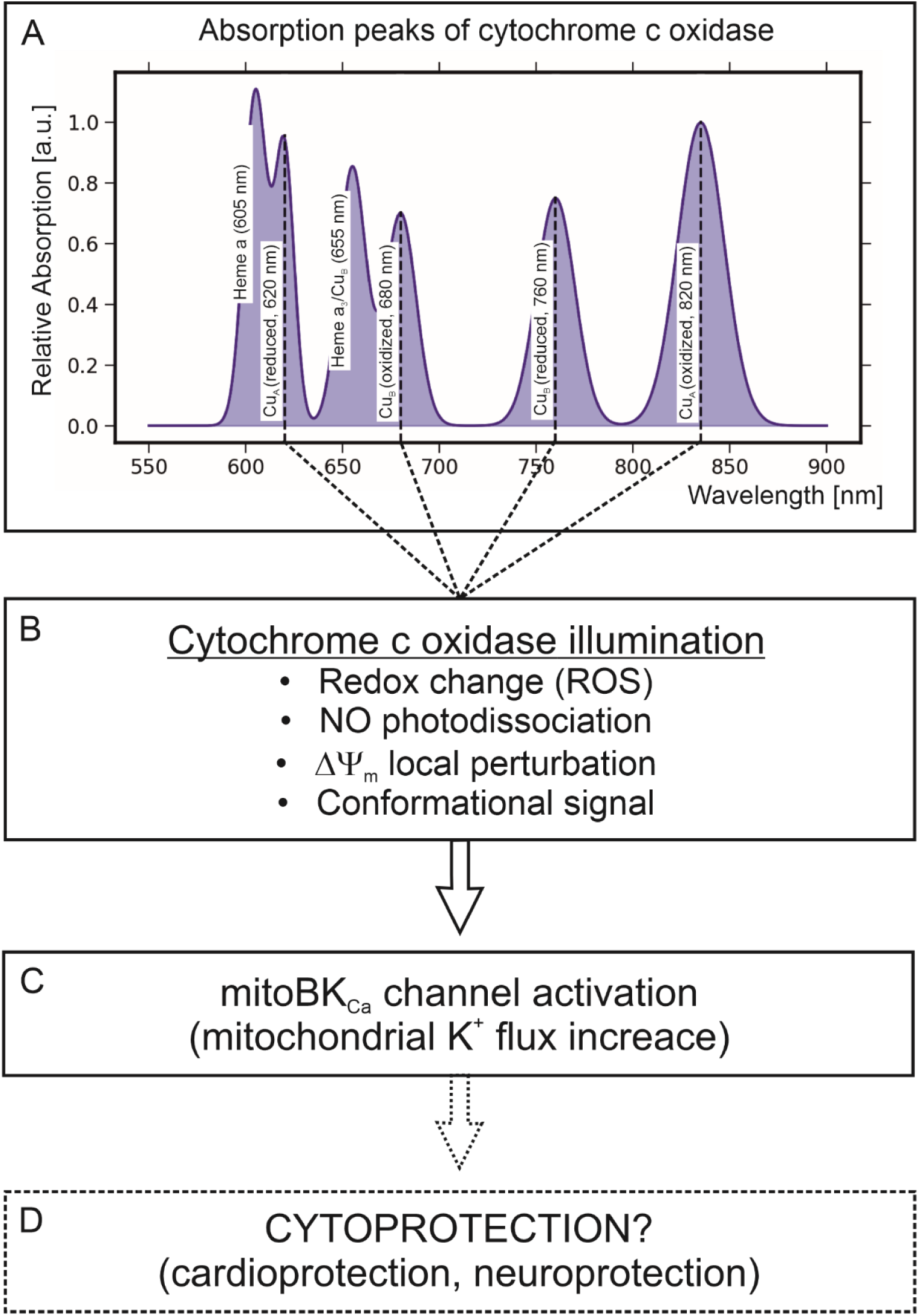
Putative mechanism of red/near-infrared light-induced activation of mitoBK_Ca_ channel. **A.** The panel shows the absorption spectrum of cytochrome c oxidase in the red and near-infrared range. Distinct absorption peaks are attributed to specific redox centers of COX, including heme a, heme a₃/Cu_B_, and Cu_A_, in either reduced or oxidized states. Labeled peaks: 620 nm, 680 nm, 760 nm, and 820 nm, indicating wavelengths at which COX can efficiently absorb light. **B.** Illumination at these wavelengths is proposed to induce functional changes in cytochrome c oxidase, including: redox modulation and reactive oxygen species (ROS) signaling, photodissociation of nitric oxide (NO), local perturbations of the mitochondrial membrane potential (ΔΨ_m_), and conformational changes in the enzyme. **C.** These mitochondrial signals are hypothesized to lead to activation of the mitochondrial large-conductance calcium-activated K⁺ (mitoBK_Ca_) channel, resulting in an increase in mitochondrial K⁺ flux. **D.** The downstream consequence of these events is proposed cytoprotection, which may contribute to cardioprotection and neuroprotection.

Future studies should aim to define the molecular determinants of the COX-mitoBK_Ca_ interaction and to determine whether the modulation involves direct redox coupling, local changes in membrane potential, or conformational signaling within multimeric protein complexes of the inner mitochondrial membrane. In addition, investigating how light intensity, exposure duration, influence channel gating will be essential for establishing quantitative relationships between light energy input and mitochondrial potassium channel responses. One should also explore the possibility that this kind of light regulation may concern other potassium channels described in inner mitochondrial membranes, such as mitoK_ATP_, mitoKv, or mitoTASK channels. Finally, the demonstration that a mitoBK_Ca_ channel can respond to light through COX-mediated mechanisms underlines the mitochondria as photoregulated organelles. This hypothesis introduces exciting possibilities for non-pharmacological modulation of mitochondrial physiology, offering new conceptual and practical avenues for developing light-based therapeutic strategies that aim to enhance cellular survival.

## ACKNOWLEDGMENTS

Supported by the Polish National Science Centre (MAESTRO grant No. 2019/34/A/NZ1/00352).

